## Supplementary Figures for "Regularized sequence-context mutational trees capture variation in mutation rates across the human genome"

### Supplemental Figures

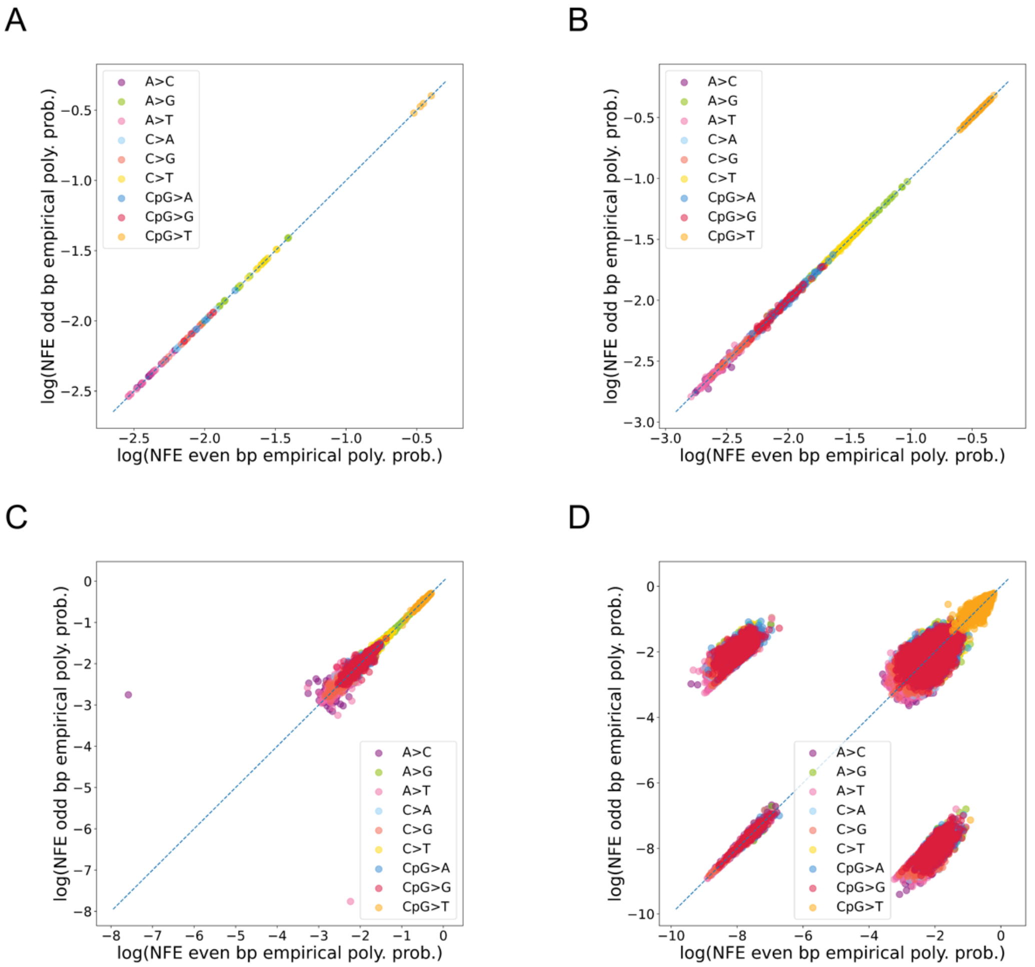


S1 Fig. Empirical even odd polymorphism probability scatter plots for the NFE dataset including zero mut variants. Baymer mean posterior estimates for (A) 3-mer models (Spearman correlation 0.999; $p < 10^{-100}$; RMSPE = 0.0009), (B) 5-mer models (Spearman correlation 0.999; $p < 10^{-100}$; RMSPE = 0.0063), (C) 7-mer models (Spearman correlation 0.992; $p < 10^{-100}$; RMSPE = 0.0459), and (D) 9-mer models (Spearman correlation 0.876; $p < 10^{-100}$; RMSPE = 0.7441) in even and odd bp datasets. Polymorphism probabilities in the bottom two, and top left, quadrants correspond to those contexts where no mutations are present for the given mutation type in the respective datasets. These polymorphism probabilities are exclusively calculated using pseudocounts.

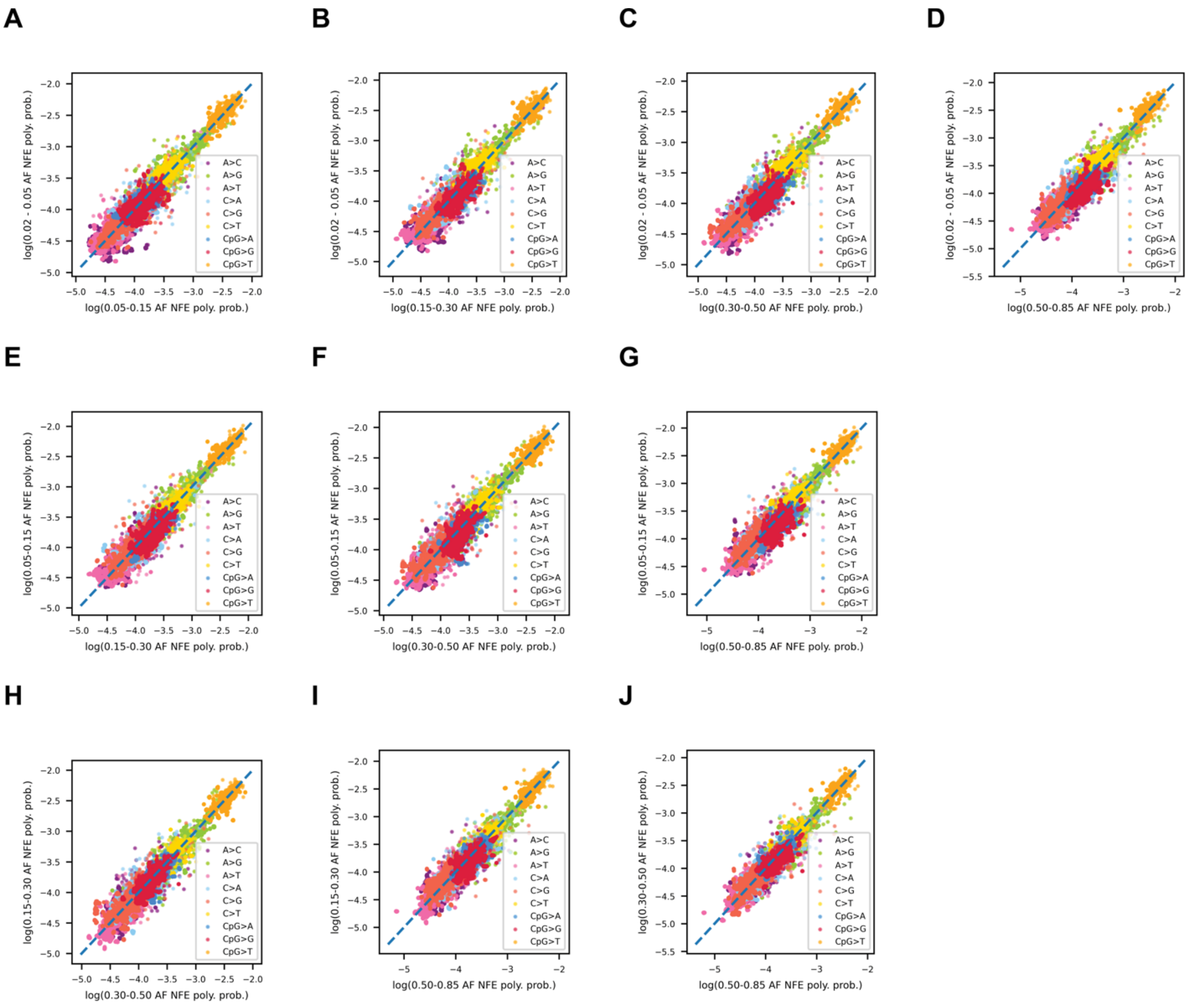


S2 Fig. Comparison of Baymer mean posterior estimates for differing allele frequency (AF) bins in the NFE dataset. (A-D) AF 0.02-0.05 compared against 0.05-0.15 AF, 0.15-0.30 AF, 0.30-0.50 AF, and 0.50-0.85 AF, respectively. (E-G) AF 0.05-0.15 AF compared against 0.15-0.30 AF, 0.30-0.50 AF, and 0.50-0.85 AF, respectively. (H-I) 0.15-0.30 AF compared against 0.30-0.50 AF and 0.50-0.85 AF, respectively. (J) 0.30-0.50 AF compared against 0.50-0.85 AF.

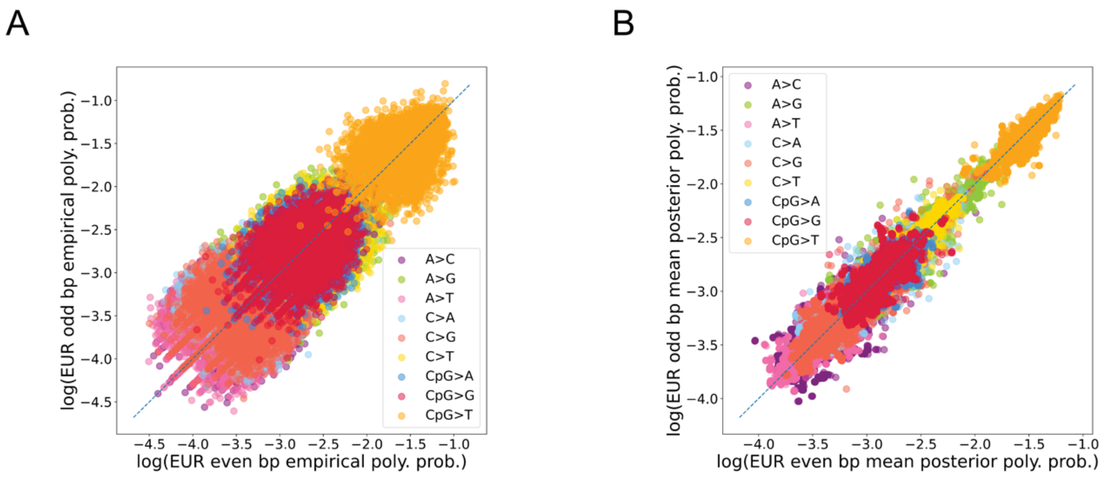


S3 Fig. Comparison of empirical and Baymer-derived 9-mer polymorphism probabilities in NYGC-resequenced 1000 Genomes Phase III non-admixed non-Finnish European (EUR) polymorphisms with derived AC >= 2 in non-coding accessible regions. (A) Empirical 9mer polymorphism probabilities for context mutations with at least 1 occurrence in both datasets (102,875 omitted context mutations) are plotted against one another (Spearman correlation 0.862; RMSPE = 0.175). (B) Baymer mean posterior estimates for 9mer polymorphism estimates in even and odd bp datasets (Spearman correlation 0.986; RMSPE = 0.042).

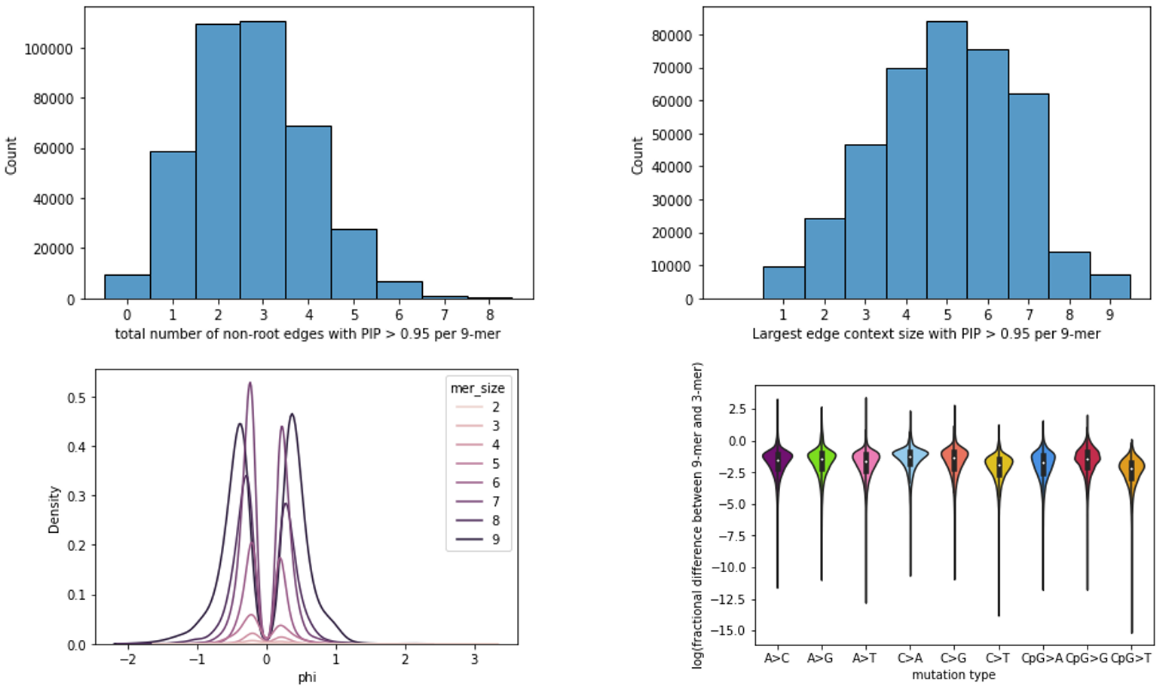


S4 Fig. Overview of the characteristics of edge mutability change dynamics in Baymer models of the NFE dataset. (A) Histogram of the number of edges per 9-mer that were inferred to confidently change polymorphism probabilities (PIP > 0.95). (B) Histogram of the maximum edge size per each 9-mer that was inferred to confidently change polymorphism probabilities (PIP > 0.95). (C) Estimated distributions of phi for each mer size level. (D) The distribution of the fractional differences of each 9-mer mean posterior polymorphism probability with their respective nested 3-mer mean posterior polymorphism probability estimates, partitioned by mutation type.

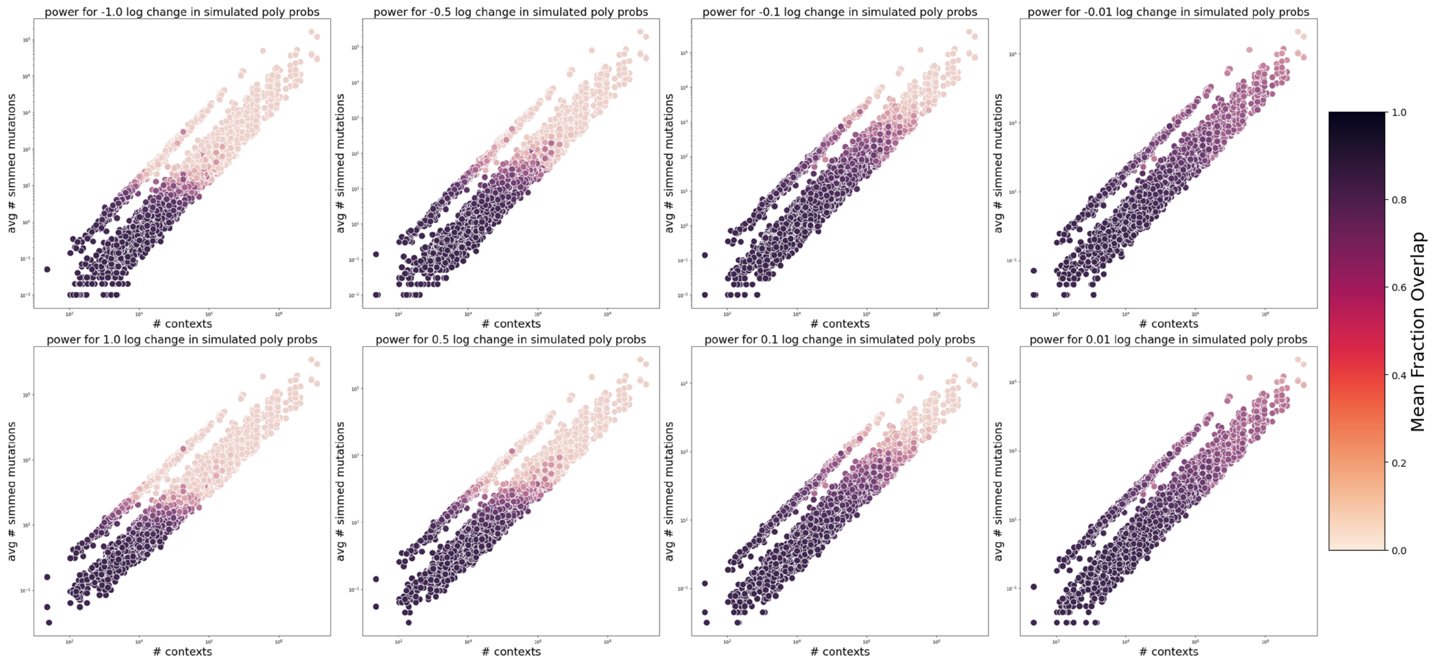


S5 Fig. Fraction overlap of simulated datasets trained by Baymer at varying sequence contexts and log changes to the null polymorphism probability.

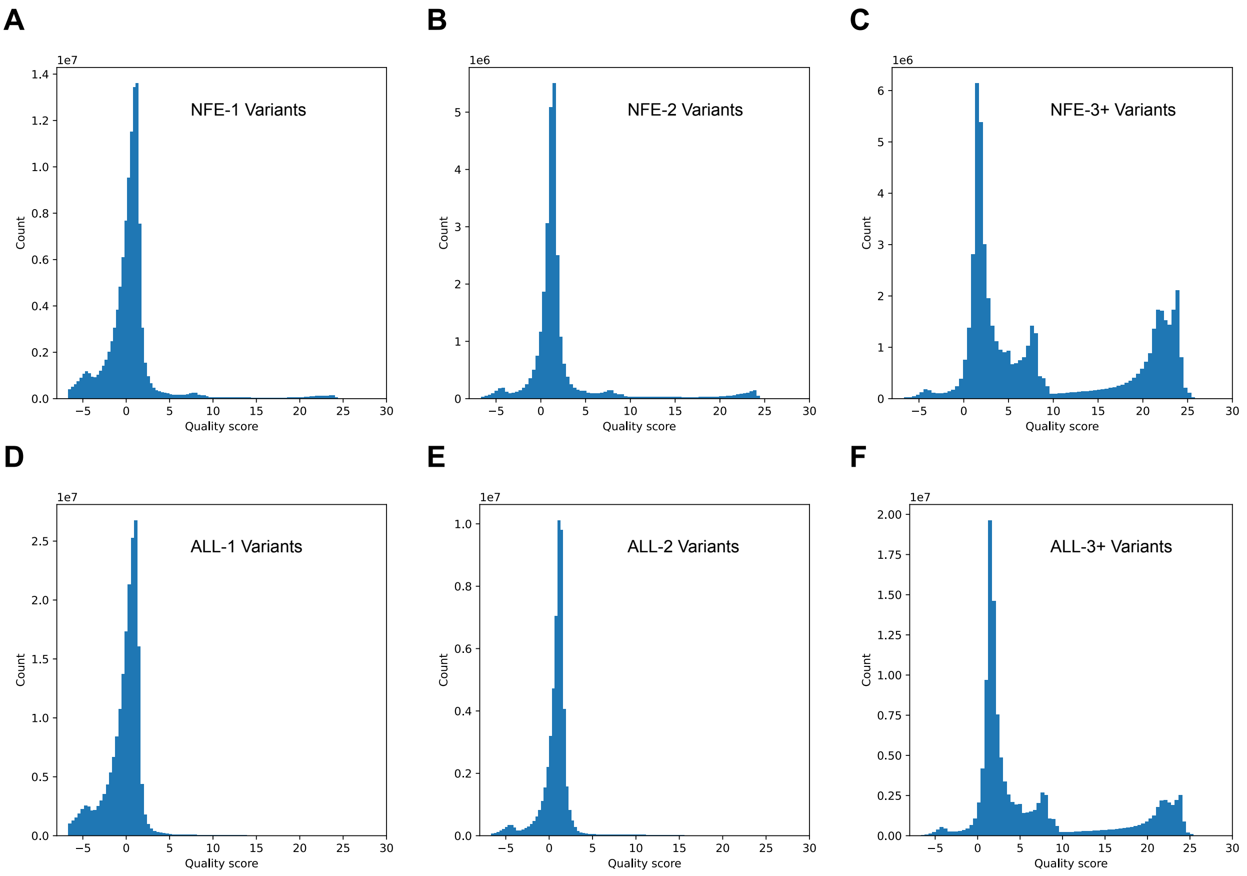


S6 Fig. Distribution of gnomAD AS_VQSLOD quality scores in non-Finnish European samples ("NFE"; A-C) and in all populations ("ALL"; D-F), separated into singletons (A,D), doubletons (B,E), and variants with allele count greater than or equal to 3 (C,F).

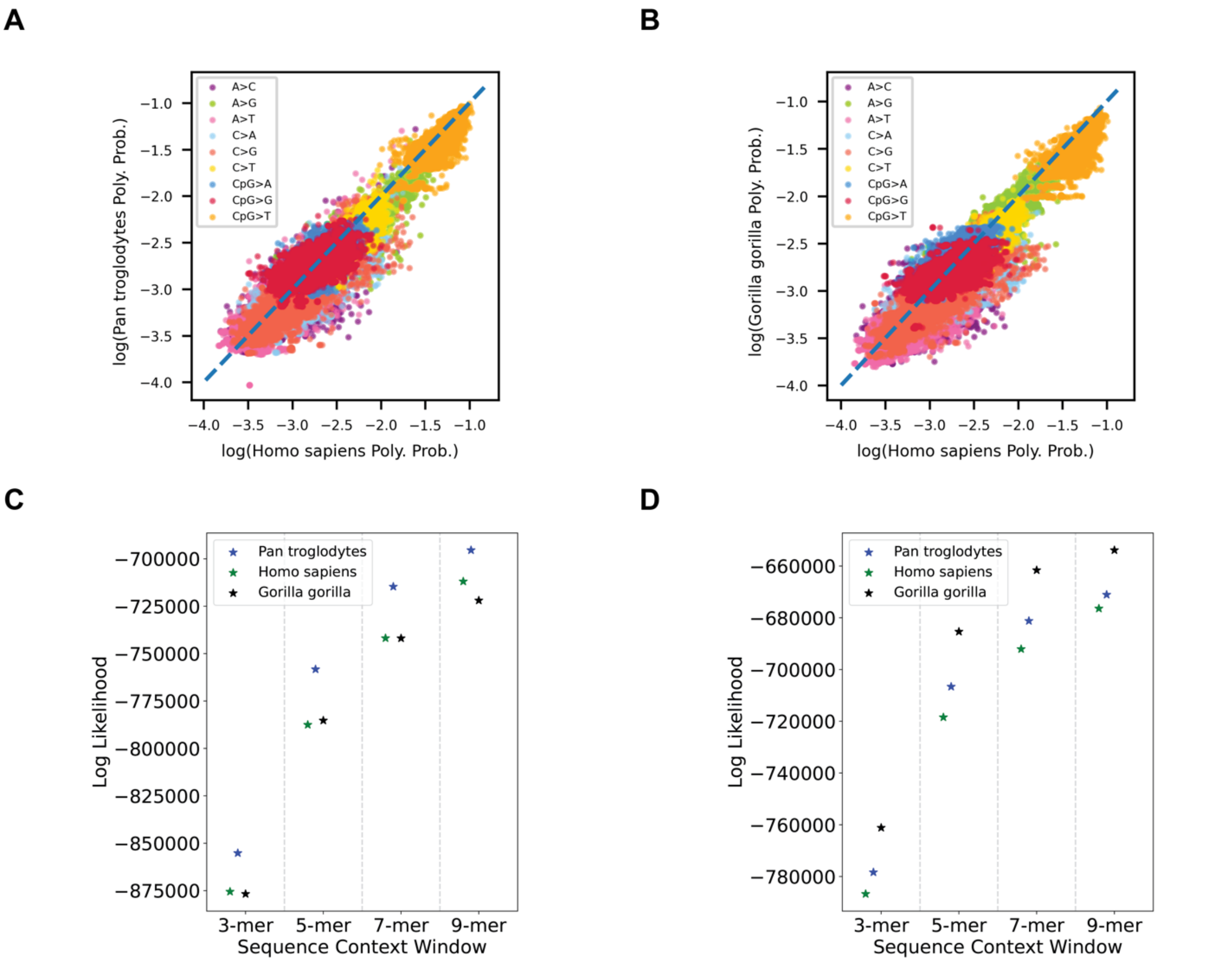


S7 Fig. Comparison of *Homo sapiens* Baymer model (NFE-2+ model) estimates with *Pan troglodytes* and *Gorilla gorilla* great ape species. (A) Mean polymorphism estimates of *Homo sapiens* model plotted against mean polymorphism estimates of *Pan troglodytes* model (Spearman correlation 0.957; RMSPE = 0.088). (B) Mean polymorphism estimates of *Homo sapiens* model plotted against mean polymorphism estimates of *Gorilla gorilla* model (Spearman correlation 0.950; RMSPE = 0.097). (C) Multinomial likelihoods for each model are calculated on odd bp pan troglodytes test data at various sequence context sizes. *Pan troglodytes* model is trained on even bp data only. (D) Multinomial likelihoods for each model are calculated on odd bp gorilla gorilla test data at various sequence context sizes. *Gorilla gorilla* model is trained on even bp data only. Polymorphism probability estimates were linearly scaled to match the mean polymorphism probability of the holdout dataset.
