## Supplementary material for "Regularized sequence-context mutational trees capture variation in mutation rates across the human genome": S1 Text

### **S1 Text - Methods**

**Sample Data Sources**

We sourced human samples from the 1KG Phase III New York Genome Center resequencing project^33^, gnomADv3.0^31^, and trios from Halldorsson et al^36^. *Pan troglodytes* and *Gorilla gorilla* samples were sourced from the Great Ape Genome Project^42^ and lifted over to GRCh38. The genomic area for all sample sources was condensed to only include coordinates included within the 1KG accessibility mask^28^ and outside of RefSeq coding regions to approximate the mappable non-coding genome. Great ape samples were additionally trimmed to remove uncallable^42^ and repeat-masked^47^ sites. Only non-indel SNVs designated as “PASS” by the data source were retained. Based on confidence calls within the FASTA sequence files, high-confidence ancestral states (designated as those sites where all sequences agree on ancestral state) were inferred for all variants and contexts within the genomic area specified, where data allowed. Otherwise, variants and sites were omitted^39^. Ancestral allele counts were used for partitioning variants into different count brackets. Variants with allele frequency greater than 0.85 were removed to control for ancestral state misidentification^48^. We also compiled all sites where the high-confidence ancestral state and GRCh38 reference genome disagree, treating this collection as a call-set of derived variants. See **S3 Text** section for URLs for all data sources.

**Baymer Model Description**

In Baymer, increasing windows of sequence context are modeled as nested trees where each sequence context has 4 children – one for each of the four nucleotides added to expand the window size. For even-sized contexts, nucleotides are added to the 5′ end, and for odd-sized contexts, to the 3′ end. In this way, sequence context trees can be iteratively constructed to a given window size. We build one such tree for every reverse-complement folded 1-mer mutation type (i.e., A>C, A>G, A>T, C>A, C>G, C>T). Note that we underline the polymorphic nucleotide in focus. For a given mutation type tree, *m*, let every edge be parameterized by $\phi_{a,b}^{m}$where *a* denotes the edge’s tree level and *b* the edge index. Edges in the first level of the tree represent the baseline A>* and C>* polymorphism probabilities (i.e., ‘1-mer’) and center the polymorphism probabilities. These edges can take any value between zero and one and are given uninformative priors $\phi_{1,0}^{m}$~ *Uniform*(0,1). All edges beyond the first levels represent the log-transformed multiplicative shifts in polymorphism probability from their respective parent nodes. The polymorphism probability for any node is therefore given by the product of the edge log-transformed multiplicative shifts leading to that node and the root node in the tree corresponding to mutation type *m*.

$p_{a,b}^{m}=\phi_{1,0}^{m}\prod_{a*,b*} exp(\phi_{a*,b*}^{m}) ;0<p_{a,b}^{m}<1$	(1)

where *a** and *b** represent the level and index of exclusively those edges leading to the context in question. For every leaf context, *I,* where the mer-level, *a*, is equal to the maximum sequence context size considered, we let ***p_i_*** denote the multinomial probabilities. Stated more explicitly:

$\boldsymbol{p}_{i}{=[p}_{a,b}^{m_{1}},p_{a,b}^{m_{2}},p_{a,b}^{m_{3}},1-\sum_{m*} p_{a,b}^{m*}] ; \sum_{m*} p_{a,b}^{m*}<1$	(2)

where *m_1-3_* denote the three mutation types possible for this context. The corresponding outcomes, ***x_i_***, for these probabilities is a length four vector for each of the three mutation types and the number of non-polymorphic context sites. We let *n_i_* denote the total number of occurrences of leaf context *i* in the genomic area specified. Over *k* leaf nodes, the likelihood for the model can be calculated as:

$p\left( y | \boldsymbol{\phi} \right)= \prod_{i}^{k} Multinom(n_{i},\boldsymbol{p}_{\boldsymbol{i}},\boldsymbol{x}_{\boldsymbol{i}})$	(3)

where *y* is a vector corresponding to the mutation counts for each mutation type and the number of unmutated loci in the given context. To provide regularization for the edges that are included in the model, we placed a spike-and-slab^29^ prior on $\phi$^22^:

$\phi_{a,b}^{m} \sim\left\{ \begin{aligned} N\left( 0,c^{2}\sigma_{a}^{2} \right) w.p. 1-\alpha_{a} \\ N\left( 0, \sigma_{a}^{2} \right) w.p. \alpha_{a} \end{aligned} \right.$	(4)

where α_a_ is the mixture probability that a given edge in mer level *a* belongs to the spike or slab. We use an uninformative prior for α_a_ ~ *Uniform*(0, 1). Both the slab and spike distribution are specified to be Gaussian with a hyperparameter, *c*, representing the ratio between each distribution’s standard deviation. The variance of the slab distribution for each level, $\sigma_{a}^{2}$, is a prespecified hyperparameter. For our models, we set this variance to ensure that the slab is favored when the evidence suggests a shift greater than 10% for a given context level (c = 500; $\sigma_{a}^{2}$ = 0.729). These chosen hyperparameters were informed by our prior biological intuition for meaningful effect sizes and a balanced ratio between the spike and slab distributions. These hyperparameters are at the discretion of the user, but a value of *c* less than or equal to 10000 is recommended^49^.

Finally, we define a latent variable, *I*, that specifies whether a given edge belongs to the spike (I=0) or slab distribution (I=1). This yields the joint posterior distribution of the model:

$p\left( \boldsymbol{\phi, I, \alpha,}\boldsymbol{\sigma}^{\boldsymbol{2}} | y \right)\propto p\left( y | \phi\right)p\left( \phi| I,\sigma^{2} \right)p\left( I | \alpha\right)p\left( \alpha\right)p\left( \sigma^{2} \right)$	(5)

To estimate the posterior distribution above, we use an adaptive Metropolis-within-Gibbs MCMC sampling scheme^30^. Every level of the tree is estimated in ascending order, setting: I=0; $\phi_{a^{'},*}^{m}$= 0 for higher-order levels (i.e., larger windows of sequence context) to aid convergence and enforce intermediate nodes to have identifiable polymorphism probabilities.

Our MCMC sampling scheme follows this approach. For the level-by-level sampling scheme, edges in levels higher, *a*′, than the level currently being sampled, *a*, are set to have no impact on the ultimate probabilities estimated, i.e., $\phi_{a^{'},*}^{m}$= 0.

For the first layer of the tree:

1. Initialize all $\phi_{1,0}^{m}$ with a random value drawn from $Uniform(0,1)$ for iteration x = 0.
2. Sample new values of each $\phi_{1,0, x}^{m}$ for this iteration *x*, from $Normal(\phi_{1,0,x-1}^{m}$, $\tau_{1,0,x-1}^{m}$) using a Metropolis step^50^, where $\tau_{1,0,x-1}^{m}$ represents the variance of the normal proposal density for $\phi_{1,0,x-1}^{m}$ at the previous iteration *x-1*.
3. Repeat step 2 until algorithm convergence.

For each subsequent level, *a* > 1:

1. Draw initial values (x=0) for parameters $\boldsymbol{\phi}_{\boldsymbol{a,b}}^{\boldsymbol{m}}$, $\boldsymbol{I}_{\boldsymbol{a,b}}^{\boldsymbol{m}}$, $\alpha_{a}$.

a. $\boldsymbol{\phi}_{\boldsymbol{a,b}}^{\boldsymbol{m}}$ is drawn from $Uniform(-0.7,0.7)$, such that the total multinomial probabilities sum to 1.

b. $\boldsymbol{I}_{\boldsymbol{a,b}}^{\boldsymbol{m}}$ is drawn from $Bernoulli(0.5)$.

c. $\alpha_{a}$is drawn from $Uniform(0,1)$.

2. Sample new values of $\boldsymbol{p}_{\boldsymbol{a,b}}^{\boldsymbol{m-1}}$ from distribution estimated in previous layer, such that the total multinomial probabilities sum to 1 given the current values of $\phi_{a,b}^{m}$.

3. Sample new values of $\phi_{a,b,x}^{m}$ from $Normal(\phi_{a,b,x-1}^{m}$, $\tau_{a,b,x-1}^{m}$) using a Metropolis step.

4. Sample new values of $I_{a,b,x}^{m}$ using a Gibbs sampling step:

$I_{a,b,x}^{m} \sim Bernoulli\left( \frac{p(I=1|\phi_{a,b,x}^{m},\sigma_{a,x},\alpha_{a,x})}{p(I=1|\phi_{a,b,x}^{m},\sigma_{a,x},\alpha_{a,x})+p(I=0|\phi_{a,b,x}^{m},\sigma_{a,x},\alpha_{a,x})} \right)$	(6)

5. Sample new values of α_a_ using a Gibbs sampling step,

$\alpha_{a,x} \sim Beta(1+\sum_{m,i=1}^{j} I_{a,b,x}^{m}, 1+j-\sum_{m,i=1}^{j} I_{a,b,x}^{m})$	(7)

where *j* represents the total number of edges in the current level.

6. Repeat steps 2-5 until algorithm convergence.

**Comparing alternate hierarchical sequence context tree architectures**

We formulated alternate trees to capture different possible relationships between sequence contexts. The asymmetric “left” alternation model is the same formulation as the default, asymmetric “right” alternation model tree, except by swapping the nucleotide addition side (5’ rather than 3’) for odd-length contexts. Nucleotides are always added to the side resulting in equal-length flanks for even-length contexts. Otherwise, the sampling and estimation scheme is identical, except with different input data for each node. To estimate symmetric models using only odd-length contexts, levels with even-length contexts were omitted, thus resulting in 16 children per node, rather than 4 in the asymmetric set-up. Model sampling and estimation are otherwise identical.

All non-Finnish European (NFE) variants with a derived allele count greater than or equal to 2 in the filtered gnomAD dataset were collected. Variants were next partitioned according to genomic coordinate parity (even/odd base pairs) to evenly divide the two groups as randomly as possible. Baymer was run on the even data sets for each tree architecture. Mean posterior estimates of polymorphism probability parameters were used as point estimates to calculate multinomial likelihoods on the odd base pair holdout data.

**Posterior coverage estimation simulations**

Polymorphism probabilities for our simulations were set using the mean of the posterior distribution estimated with Baymer when applied to private European variant data with minimal jitter added to avoid over-regularized estimates while still maintaining realistic human context-dependent polymorphism probability patterns. Jitter was added by sampling every 9-mer polymorphism probability, $p_{a,b}^{m}$, from $Normal(p_{a,b}^{m}, {{(p}_{a,b}^{m})}^{1.5}$), where the variance was set to scale to the underlying polymorphism probability. This dataset was chosen as it had the property of reaching sparsity limits at the 7-mer level and beyond. Thus, simulations evaluated up to 7-mers would provide a mixture of sparse and data-rich sequence contexts, providing a representative proxy for larger datasets run up through the 9-mer level. Using these polymorphism probabilities, new datasets were simulated by sampling from the multinomial distribution for each 9-mer sequence context. After applying Baymer to each individual dataset, we calculated the frequency that the true polymorphism probabilities were included in different sized credible sets. 2000 simulations were run for every sequence context up until 7-mers. Equal-tailed intervals were used to assign the credible intervals. Note that to aid computational tractability of this number of contexts and simulations, the alpha mixing parameter was sampled by using the posterior distributions for each level of the underlying base probability model used to generate simulated data.

**Model comparisons for even/odd base-pair subsets**

The same NFE even/odd variant partitions were used as in the alternation pattern experiments above. Baymer was run on even and odd sets independently and the mean posterior estimates of polymorphism probability parameters were returned.

The root mean squared perpendicular error (RMSPE) was calculated by measuring the perpendicular distance between each point (estimated polymorphism probability) and the x=y line, that assumes each estimate is identical between models.

For transferability experiments, all European samples, excluding Finnish samples, from the 1KG Phase III designated as non-admixed^28^ were aggregated and trimmed to only include sites with a minimum of 2 derived alleles and again partitioned according to genomic position parity. Opposite parities between 1KG and gnomAD datasets were grouped together. For each dataset, 100 equally sized allele frequency bins between the minimum allele frequency in the two datasets and 1.0 were set. Each dataset was randomly downsampled to ensure the same number of variants in each allele frequency bin. Baymer was applied to each down-sampled dataset and mean posterior estimates were compared.

**Extended Sequence Context Likelihood Estimation**

The gnomAD NFE data was partitioned into even and odd base pairs as described above. For each split, models were estimated using Baymer up through 9-mers. Smaller models correspond to the Baymer tree with all edges in larger sequence contexts not being considered assigned uninformative shifts ($\phi_{a,b}^{m}=0$). We calculated likelihoods using the mean posterior probability estimate at the 9-mer level on the opposite parity polymorphism count data.

**Data Sparsity Filters**

To distinguish the degree to which estimates of PIP are simply a byproduct of data sparsity, we filtered out all sequence contexts with fewer than 50,000 total instances or fewer than 50 mutations in the non-coding genomic area considered.

**Identification of Sequence Context Motifs**

We examined the 100 most and least mutable sequence contexts for each mutation type in the NFE model and manually identified recurrent patterns in these contexts, with a slight preference towards longer motifs. We identified 66 total motifs and performed a hypergeometric enrichment test for the top and bottom 1% of contexts based on polymorphism probability. We report Bonferroni-corrected p-values for each context initially chosen (66 tests).

**Private Variant Analyses**

All continental populations without substantial recent admixture (African, European, South Asian, East Asian) from the NYGC 1KG phase III resequencing dataset were filtered to only include variants private to each continental group. Each population was trimmed to only include variants with a minimum allele count of 2 and then down-sampled and site frequency spectra-matched to match the smallest variant counts across the four continental groups. Baymer was then applied to each resulting dataset. The resulting posterior distributions of the polymorphism probabilities and $\phi$ shifts of each model were then pairwise compared by calculating the fraction overlap of the distributions, as a proxy for the probability they are the same. Distributions are parameterized using a Gaussian kernel density estimate on the posterior samples.

**Power Estimates**

Truth polymorphism probabilities used in our simulations to estimate power were set using the same model as the variance calibration experiments. For a given sequence context mutation, we tested the discoverability of a spectrum of deviations from the “truth” model. We simulated 1000 9-mer count tables using polymorphism probabilities from both the “truth” model and the deviated model. Both count tables were modeled using Baymer and the resulting posterior distributions used to assess the fraction overlap for the context mutation in focus. A shift is considered discovered if the degree of fraction overlap is less than 1%. As running this experiment for all context mutations was intractable, we tested at most 100 CpG and 100 non-CpG contexts at each mer-level. Contexts were chosen to give an even spread across the sample size spectrum, as dictated by total contexts.

**Grafted Tree Scheme**

Baymer models were built independently on *de novo* even data and gnomAD NFE polymorphism data with allele count greater than or equal to 2. The *de novo* model parameter estimates were used up through 3-mers. For the remaining levels (for 5-mers and larger windows), NFE-2+ parameter mean point estimates were used in place of the equivalent de novo edges. Thus, the grafted tree polymorphism parameters were the product of the point estimates for each branch of the tree, given the data source described above. The multinomial likelihood of the resulting model was calculated on the odd de novo holdout data, as before.

**Great Ape Model Comparisons**

*Pan troglodytes* and *Gorilla gorilla* datasets were modeled using Baymer and mean posterior estimates of 9-mer polymorphism estimates were compared with equivalent estimates from the NFE-2+ model scaled to match each comparison model’s total polymorphism probability. For likelihood calculations, the focal population (here *Pan troglodytes* and *Gorilla gorilla*, respectively) was divided into even and odd base pair data, using the odd base pair data as the test set and even base pair data as the training set. For other populations, the entire dataset is used to calculate likelihoods after adjusting each comparison model with a scalar to normalize for the total polymorphism probability of the test set.
