## Supplementary material for "Regularized sequence-context mutational trees capture variation in mutation rates across the human genome": S3 Text

### **S3 Text – Data Availability**

We have implemented our Baymer method into software that is freely available as a python package. This can be accessed on the Voight Lab GitHub repository: <https://github.com/bvoightlab/Baymer/>. Additional outputs generated by the model presented in this work are also available at: <https://doi.org/10.5281/zenodo.7843023>. All data analyzed here are publicly available at the following websites:

**NYGC resequencing of 1KG Phase III data:**

http://ftp.1000genomes.ebi.ac.uk/vol1/ftp/data_collections/1000G_2504_high_coverage/working/20190425_NYGC_GATK/

**gnomADv3.0:**

https://gnomad.broadinstitute.org/downloads

**Halldorsson et al. trio data:**

https://science.sciencemag.org/highwire/filestream/721792/field_highwire_adjunct_files/7/aau1043_DataS5_revision1.tsv

**1KG accessibility mask:**

http://ftp.1000genomes.ebi.ac.uk/vol1/ftp/data_collections/1000_genomes_project/working/20160622_genome_mask_GRCh38/PilotMask/20160622.allChr.pilot_mask.bed

**RefSeq coding regions:**

http://www.ensembl.org/biomart/

**Ancestral FASTA:**

<ftp://ftp.ensembl.org/pub/release97/fasta/ancestral_alleles/homo_sapiens_ancestor_GRCh38.tar.gz>

**Great Ape Genome Project Data:**

https://www.biologiaevolutiva.org/greatape/data.html
