## Supplementary material for "Regularized sequence-context mutational trees capture variation in mutation rates across the human genome": S2 Text

### **S2 Text – Computational Considerations**

Baymer can estimate a full 9-mer model using 8 threads and ~12 GB of RAM in ~5.5 hours. There are no inherent model constraints on the size of the sequence context window estimated outside of computational complexity. Given the trajectory of models run using 8 threads for smaller context window sizes, we expect approximately 4x additional time for each context expansion (**S8 Table**), consistent with the increase in total parameters.
